# Bin2cell reconstructs cells from high resolution Visium HD data

**DOI:** 10.1101/2024.06.19.599766

**Authors:** Krzysztof Polański, Raquel Bartolomé-Casado, Ioannis Sarropoulos, Chuan Xu, Nick England, Frode L. Jahnsen, Sarah A. Teichmann, Nadav Yayon

## Abstract

**Summary:** Visium HD by 10X Genomics is the first commercially available platform capable of capturing full scale transcriptomic data paired with a reference morphology image from archived FFPE blocks at sub-cellular resolution. However, aggregation of capture regions to single cells poses challenges. Bin2cell reconstructs cells from the highest resolution data (2 *μ*m bins) by leveraging morphology image segmentation and gene expression information. It is compatible with established Python single cell and spatial transcriptomics software, and operates efficiently in a matter of minutes without requiring a GPU. We demonstrate improvements in downstream analysis when using the reconstructed cells over default 8 *μ*m bins on mouse brain and human colorectal cancer data.

**Availability and Implementation:** Bin2cell is available at https://github.com/Teichlab/bin2cell, along with documentation and usage examples, and can be installed from pip. Probe design functionality is available at https://github.com/Teichlab/gene2probe

**Supplementary information:** Supplementary data are available online.

## Introduction

Recent years have seen massive advances in spatial transcriptomics, with novel platforms like Stereo-Seq and S1000 overcoming technical compromises of earlier methods and producing comprehensive gene expression information at sub-cellular resolution (Wang *et al*., 2023). Visium HD is further compatible with archived (FFPE) sample blocks, which dramatically expands its potential sample pool and holds additional promise for paired high-resolution morphological assessment (Nagendran *et al*., 2023). The array features full coding transcriptome probes placed on a gap-less grid, barcoded in 2 *μ*m square regions (bins) which are then grouped into 8 *μ*m square bins for default analysis or annotation. The data is accompanied by a matching high resolution bright field (e.g. hematoxylin & eosin, H&E) or fluorescent (e.g. immunofluorescence, IF) morphology image. While the 8 *μ*m resolution is a big improvement over original Visium’s 50 *μ*m spots, having access to 2 *μ*m bins along with matching morphology information makes it tempting to reconstruct single cells from the data.

Bin2cell operates on Visium HD’s highest 2 *μ*m resolution, joining the sub-cellular bins into single cells. This is done by performing morphological segmentation with StarDist (Schmidt *et al*., 2018), identifying nuclei with its pretrained H&E model and subsequently expanding them to neighbouring unlabelled bins. Areas where nuclei were not captured in the H&E image, or take on unusual shapes the model can’t detect, have secondary labels identified based on spatial clusters of expression count totals. The software also offers a number of utility functions, such as a data loader accounting for the new Visium HD quantification pipeline formatting, variable bin size correction of the transcriptomic data, custom resolution image creation and storage for superior plotting, and an additional workflow to create custom probes if wishing to enhance the default panel. The package is implemented with minimising RAM use and run time in mind, and is fully compatible with SCANPY (Wolf *et al*., 2018), the widespread Python single cell/spatial transcriptomics analysis standard. Bin2cell is freely available on GitHub, along with documentation and examples, and can be easily installed from pip. We demonstrate the package’s utility by favourably comparing its proposed cells to the 8 *μ*m resolution bins on mouse brain and human colorectal cancer data.

## Materials and methods

The bin2cell workflow is illustrated in Fig 1a and Supp Fig 1,2. The package is SCANPY (Wolf *et al*., 2018) compatible, unlocking a number of processing and visualisation options along with access to third party software like CellTypist (Domínguez Conde *etal*., 2022). Given the scope of Visium HD data (millions of bins by tens of thousands of genes, morphology images up to 10GB), care was put into streamlining all operations for run time and RAM use. As a result of the combined efficiency of StarDist and bin2cell, analysing the mouse brain demo data from raw input to cell object takes 15 minutes on CPU, with RAM use not exceeding 10GB.

**Fig. 1.**
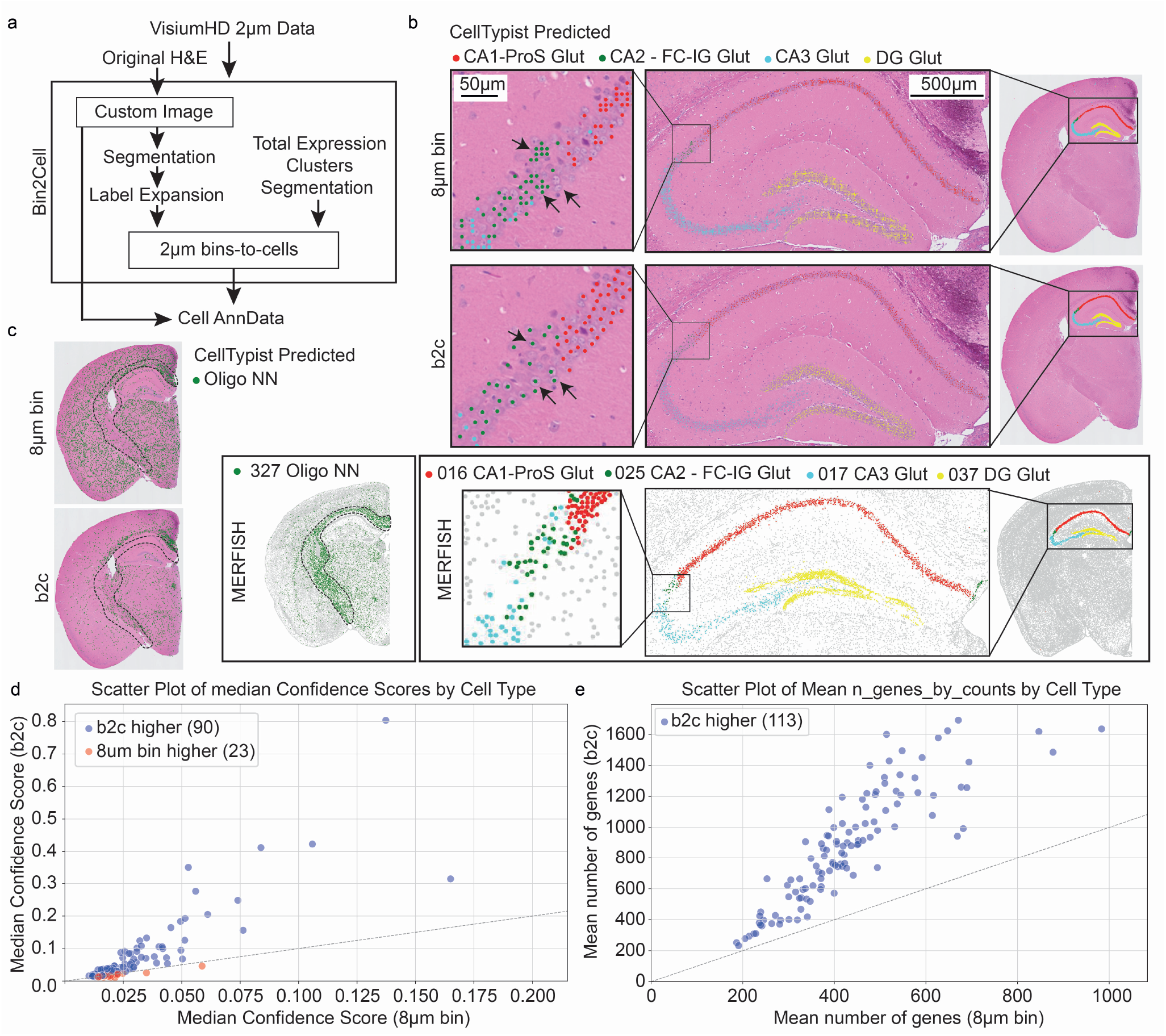
Bin2cell workflow and application to mouse brain Visium HD. **a**. Schematic illustration of the bin2cell pipeline, utilising both the original image and gene expression data to group 2 *μ*m bins to cells. The output is a standard AnnData spatial object. **b**. Comparison of 8 *μ*m (top) default Visium HD object compared to bin2cell (middle) and expert annotated (Allen brain atlas) single cell MERFISH spatial data (bottom). CellTypist predictions were derived for all cells and hippocampal formation neurons are shown for both Visium HD objects. Arrows illustrate the over/under representation of ‘cells’ by the 8 *μ*m object compared to bin2cell segmentation. CellTypist confidence score > 0.05. **c**. Comparison of CellTypist predictions for oligodendrocytes highlighting the lack of specificity of the 8 *μ*m object. CellTypist confidence score > 0.01. **d**. CellTypist model median confidence scores for all cell types, a minimum of 0.01 median confidence score for b2c and 8 *μ*m object, blue are cell types that have a higher score for b2c. **e**. Mean number of genes in cell types in b2c compared to 8 *μ*m, all cell types are higher in b2c

### Custom resolution H&E image

The tissue image created by Space Ranger (10X’s Visium HD mapping and quantification software) is scaled to a maximum dimension of 6000 pixels, rendering it unfit for segmentation due to insufficient resolution. In turn, processing the full scale H&E image runs the risk of objects being too large to detect in the case of high zooms, in addition to expending unnecessary resources processing areas of the tissue image not part of the capture region. An early step of bin2cell creates a controlled resolution reference image limited to the capture region, ensuring consistent and reproducible input for segmentation and plotting (Supp Fig 1B). The resolution is set in micrometers per pixel (mpp), with a recommended value of 0.5.

### Variable bin size correction of transcriptomic data

Visium HD has a novel technical effect – the 2 *μ*m bins come with a size disparity on a per-row/per-column basis, leading to a characteristic ‘striped’ appearance when visualising total counts per bin across the tissue. To correct for this, each bin’s count total is divided by a user-specified quantile (by default 0.99) from its corresponding array row, with the procedure subsequently repeated on a per-column basis. The resulting factors are brought back into count space by multiplying them by the specified quantile of global per-bin count totals. This greatly reduces the ‘striped’ effect, as shown in Supp Fig 1C. The count matrix is then by default rescaled per bin to match the corrected count totals.

### Morphology and gene expression image segmentation

Nuclei are identified in the H&E image by using StarDist (Schmidt *et al*., 2018), an efficient segmentation algorithm that comes with a pretrained H&E model. The nuclei labels are discretised into bins, and expanded into cells by label expansion of a fixed distance (by default 2 bins, i.e. 4 *μ*m). Unassigned bins equidistant to bins from multiple nuclei are assigned based on transcriptional similarity, as represented in PCA space.

An optional secondary segmentation is performed where the total gene expression per bin is represented on the Visium HD array grid, identifying spatial clusters via StarDist’s fluorescence model. This segmentation is less robust than the H&E nucleus detection, struggling in denser tissue regions, but can identify putative cells where the morphology failed to detect nuclei due to them not appearing in the tissue slice or taking on unusual shapes. Only labels exclusive to bins unassigned after morphological segmentation are taken for subsequent use. An example evolution of the detected labels as the workflow progresses is shown in Supp Fig 2.

The final object accumulates all counts from 2 *μ*m bins by cell labels as well as the new spatial location of cells, the custom H&E image and the number of bins that were accumulated for each cell.

### Custom probe creation

It is possible to enhance Visium HD panels with additional probes, capturing particular isoforms or sex-specific gene expression. Gene2probe finds the best candidate probes for a gene of interest by evaluating all possible sequences against the 10X Genomics probe construction specifications. More details in the supplement, with the package freely available at https://github.com/Teichlab/gene2probe.

## Results

### Bin2cell recapitulates single-cell spatial and transcriptomic profiles of the mouse brain

To test the performance of bin2cell (b2c) segmentation as opposed to the default 8 *μ*m bin object, we used demo Visium HD mouse brain data distributed by 10X Genomics. The mouse brain is perhaps the most broadly studied and annotated biological example, and as such serves as a fantastic case to test bin2cell segmentation. The most recent single-cell atlas (Yao *et al*., 2023) was used to construct a CellTypist (Domínguez Conde *et al*., 2022) model, predicting the annotations of both b2c and 8 *μ*m representations of the data. The spatial locations were subsequently compared to expert annotated, high-resolution, *∼*550 gene custom panel MERFISH data (Zhang *et al*., 2023), treated as the ground truth to judge the results against.

Broadly, both b2c and 8 *μ*m binned data recapitulated many expected cell positions and distributions across the entire brain, with b2c data more accurately depicting the distribution of cells compared to reference MERFISH data (Fig 1b-c, Supp Fig 3-5). To compare the cellular distributions between objects, we selected the hippocampal formation, with CA1, CA2, CA3 and dentate gyrus (DG) regions seemingly accurately recapitulated in both Visium HD objects on the macro scale. However, on the cellular level, 8 *μ*m bins fail to account for cell positions and either over represent cells by clusters of bins or fail to capture cells in other cases. In contrast, b2c closely follows nuclei positions and cellular distributions in the tissue due to utilising information from the morphology image (Fig 1b). The increased confidence of interpreting b2c output as cells, coupled with the improved resolution, will enable cell communication analysis in ways previously impossible.

There are cell types for which bin2cell drastically outperforms the 8 *μ*m bins in terms of inferred spatial location. An example are oligodendrocytes, where the 8 *μ*m bin CellTypist predictions appear far more often than expected in grey-matter regions. The expected white-matter locations have fewer oligodendrocyte calls than anticipated, likely due to regions of very sparse data that have few bins present in the area after a lenient 100 unique gene filter. By contrast, b2c identifies the population closer to the MERFISH ground truth, including overcoming the sparsity of the white-matter regions and proposing more *≥*100 unique gene cells there (Figure 1c, Supp Fig 3). Finally, we used the obtained CellTypist confidence scores as a measure of quality of proposed single cell information. Of the 113 cell types, 89 have median confidence scores higher in b2c than in 8 *μ*m bins (Figure 1d). All 113 cell types have a higher average number of genes detected in b2c (Figure 1e). Increased gene coverage will allow novel cell function discovery not restricted to prior knowledge or specific panel design.

While the 8 *μ*m bins seem to match expected cell positions, the ability to infer cell-to-cell association would be significantly hampered by issues stated above. To estimate this, we calculated the mean neighborhood proportions of cells in the mouse brain, reporting an average per cell type. The MERFISH data was used as a reference to compare the distributions to. Bin2cell neighborhood proportion patterns matched the MERFISH much better (Mean Square Error, MSE - 0.00086) compared to 8 *μ*m bins (MSE - 0.00167) (Supp Fig 6).

### Bin2cell analysis of human colorectal cancer empowers high resolution morphological inference

We also applied bin2cell to 10X Genomics demo human colorectal cancer data. The gut is a perfect tissue for 8 *μ*m binning to work well due to a high density of small cell clusters of similar cell types. Nevertheless, bin2cell segmentation retains an advantage when evaluated using the same criteria as the mouse brain. While providing modest improvements to the 8 *μ*m bins in CellTypist prediction confidence, bin2cell outperforms 8 *μ*m bins in per-cell gene count coverage and the ability to recapitulate fine morphological structures. This manifests in a superior reconstruction of venous and atrial layers, akin to the hippocampal formation from the mouse brain, as well as behaviour along the tissue edge. The 8 *μ*m object has a number of bins with errant predictions, necessitating secondary filtering. Bin2cell’s use of morphology ends up providing a bin grouping that gets correctly predicted as CMS2 (Supp Fig 7-8).

## Acknowledgments

The Wellcome Sanger Institute is supported by core funding from the Wellcome Trust (220540/Z/20/A). This work was supported by the Engineering and Physical Sciences Research Council (grant number EP/Y02978X/1).

