## Supplementary Material for "Bin2cell reconstructs cells from high resolution Visium HD data"

### Supplementary sections

#### Appendix

1) Bin2cell pipeline detailed figures
    a) Supplementary figure S1. Extended bin2cell pipeline and image preprocessing
    b) Supplementary figure S2. Bin2cell segmentation, label expansion and harmonisation.

2) Supplementary Section 2: Limitations for the 8μm bin object for single cell analysis.

3) Spatial mapping of mouse coronal hemibrain
    a) Spatial mapping of cortical cell types
    b) High resolution cell neighbourhood analysis

4) Spatial mapping of human colorectal cancer
    a) Cell type prediction by a combination of CellTypist models
    b) Prediction performance and gene coverage
    c) Spatial patterns on tissue edges and vasculature

5) Gene2Probe custom probe design for Visium CytAssist and FLEX

### Supplementary Section 1: Bin2cell pipeline detailed figures

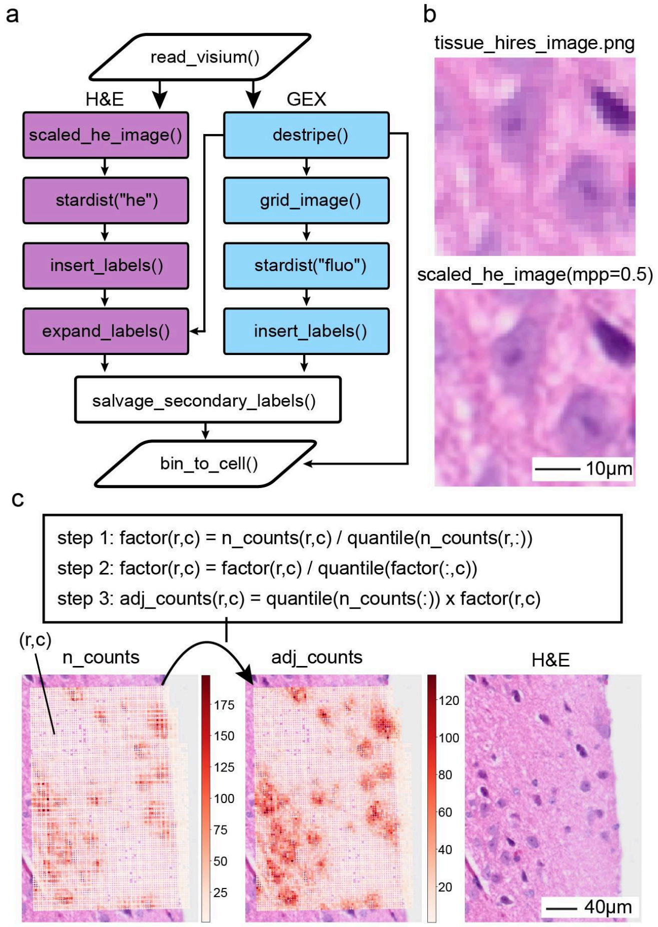


**Supplementary Figure S1. Extended bin2cell pipeline and image preprocessing. a.** Outline and functionality of bin2cell processing pipeline. Bin2cell utilises a custom Visium HD import function. Image (e.g. H&E) and gene expression data (GEX) is processed in two routes. A custom H&E image is generated based on user defined resolution as shown in (b). The image is segmented with the StarDist pretrained H&E model and generated labels are migrated to the 2μm object. Finally, H&E labels can be expended by proximity and conflict spots are resolved by gene expression similarity (see Supp Fig 2d). The capture field (or lawn) holds row and column capture variance which we can correct with an iterative algorithm described in (c). We also support the creation of a custom grid image of the corrected total counts RNA coverage. This image is segmented with the fluorescence StarDist model to capture RNA clusters and labels are imported to the 2μm object. The last step utilises the GEX labels by adding segmented GEX labels that were not captured by expanded H&E labels. **b.** Illustration of SpaceRanger highest-resolution image which is suboptimal for segmentations compared to the custom mpp (microns-per-pixel) image (bottom). **c.** Correction of row and column bias.

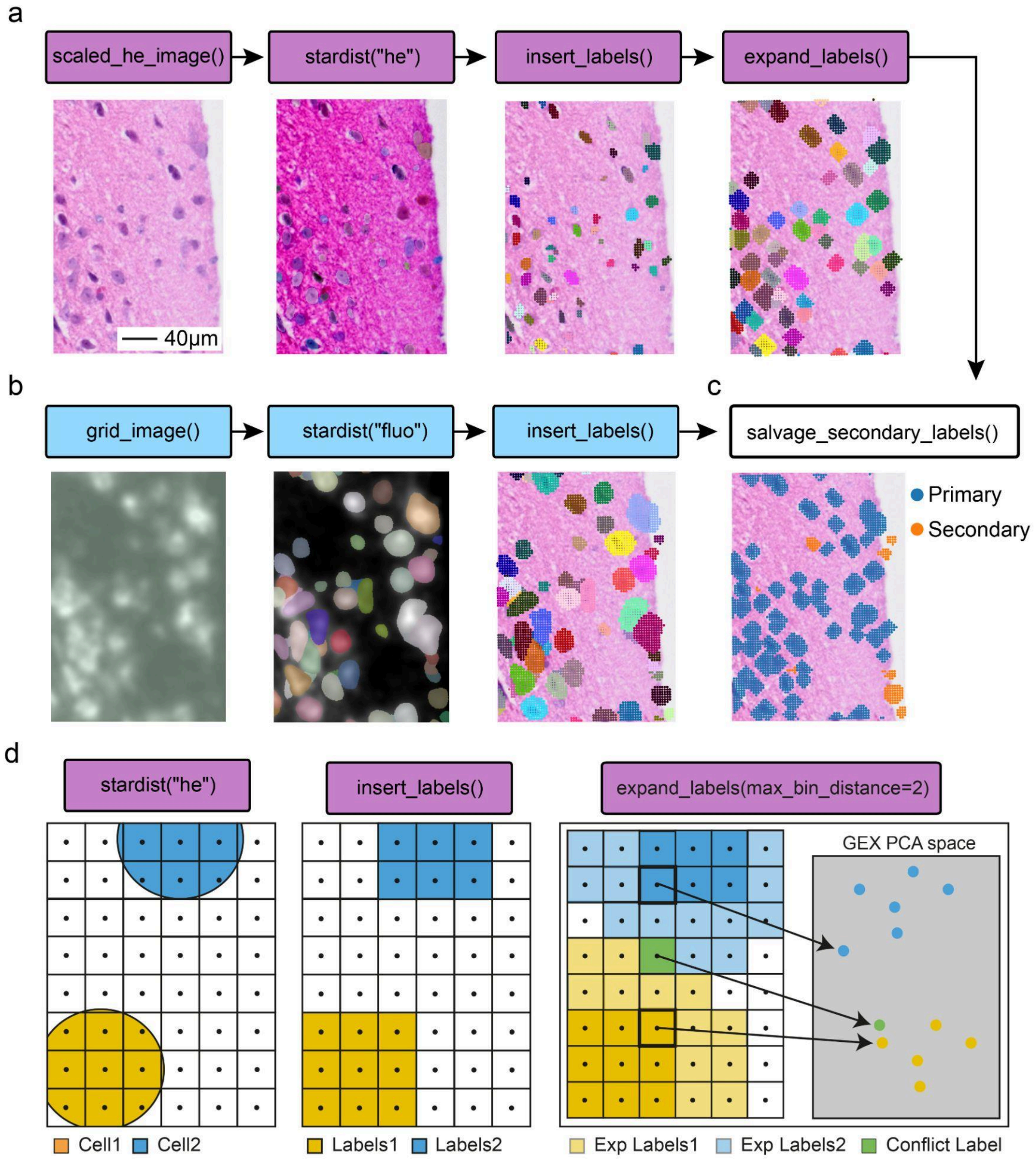


**Supplementary Figure S2. Bin2cell segmentation, label expansion and harmonisation.**
**a. Example region for H&E nuclei segmentation.** Custom scaled H&E image created by scaled_he_image() serves as the basis for large scale StarDist segmentation via stardist("he"). The predicted nuclei labels are transferred to the 2μm spatial object by insert_labels(). The original nuclear labels are expanded by adding "layers" of 2μm GEX spots. The number of layers is user specified. **b. GEX image segmentation.** grid_image() generates a mockup image based on the number of counts per spot generating clusters of RNA expression. This image is

then segmented with StarDist with the pertained fluorescent model to generate labels that are transferred to the 2µm spatial object by insert_labels(). **c.** To take advantage of both label types, GEX labels that are not found in H&E are added while overlapping GEX segmentations are ignored. **d.** Label expansion strategy. 2µm GEX spots (square grid) are assigned by spot centroid (dots) overlap to the segmented nuclear label. Additional spots are assigned to nearby cells by Euclidean distance to the external nuclei spots. Spots that are equidistant from 2 cells are assigned by GEX proximity in PCA space to the closest original cell spots.

#### Supplementary Section 2: Limitations for 8µm bins for single cell analysis.

The 8µm bin default object provided by 10X Genomics is the only resolution available for user annotation and visualisation with the Loupe browser provided for Visium HD. This representation of the data aggregates all RNA counts from sixteen adjacent 2µm bins into a single 8µm bin. While this approach appears to address data sparsity issues, applying a set **grid structure** effectively acts as a spatial filter with a predetermined frequency and shape. As such, it will highlight spatial patterns that match these specific properties and diminish others:

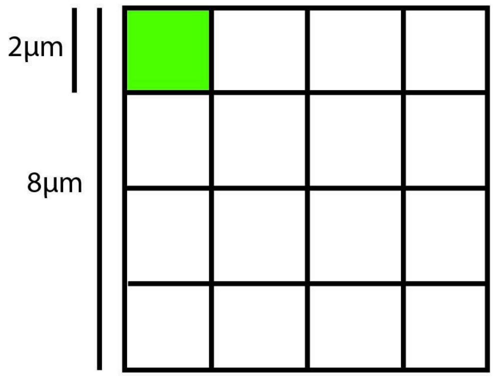


1) Large cells will be fragmented by multiple bins while small or non-square cells would have suboptimal overlap with the square 8µm bins.
2) Downstream analysis will be heavily biased by the grid properties. The typical distances between cells will be units of 8µm (e.g. 8, 16, 24 etc). This will diminish the ability to infer realistic cell associations, or cell-cell communication.
3) Boundaries, fine tissue structures (e.g. vessels) or tissue edges will be affected as these will deviate dramatically from a square grid.
4) 8µm bins are likely to capture contents of multiple cells, introducing intrinsic doublets drastically hampering the ability to explore the true molecular content of a single cell in an unbiased manner, which is a large promise for full transcriptome spatial transcriptomics.

### Supplementary Section 3: Cell type predictions in mouse brain

The “Visium HD Spatial Gene Expression Library, Mouse Brain (FFPE)” Visium HD demo dataset by 10X Genomics was retrieved from https://www.10xgenomics.com/datasets/visium-hd-cytassist-gene-expression-libraries-of-mouse-brain-he

As part of bin2cell, images were rendered at 0.5 mpp. StarDist H&E segmentation was performed with prob_thresh=0.01 and fluo segmentation was performed with prob_thresh=0.05 and nms_thresh=0.5.

The 8µm bins object from the mouse brain has 393,543 bins. In contrast, b2c finds 69,902 cells in the same section which is much closer to the expected number of cells in the mouse brain coronal hemisphere cut at 5µm. For reference, MERFISH data that was obtained from a 10 µm frozen section of 2 hemispheres has 130,112 cells. Removing cells with low gene capture (<100 genes per cell) results in 60,274 cells for b2c and 266,848 bins for the 8µm object. The expression matrices were normalised to 10,000 counts per cell/bin and log1p-transformed prior to CellTypist predictions using the “Mouse_Whole_Brain” model.

In terms of cell distributions, both objects reliably capture the expected broad cell patterns. However, for the 8 µm bins, the cell numbers in grey matter regions are inflated and cells in white matter regions are filtered out by a minimal filter of 100 genes per cell. (**Supplementary Figure s3-5**)

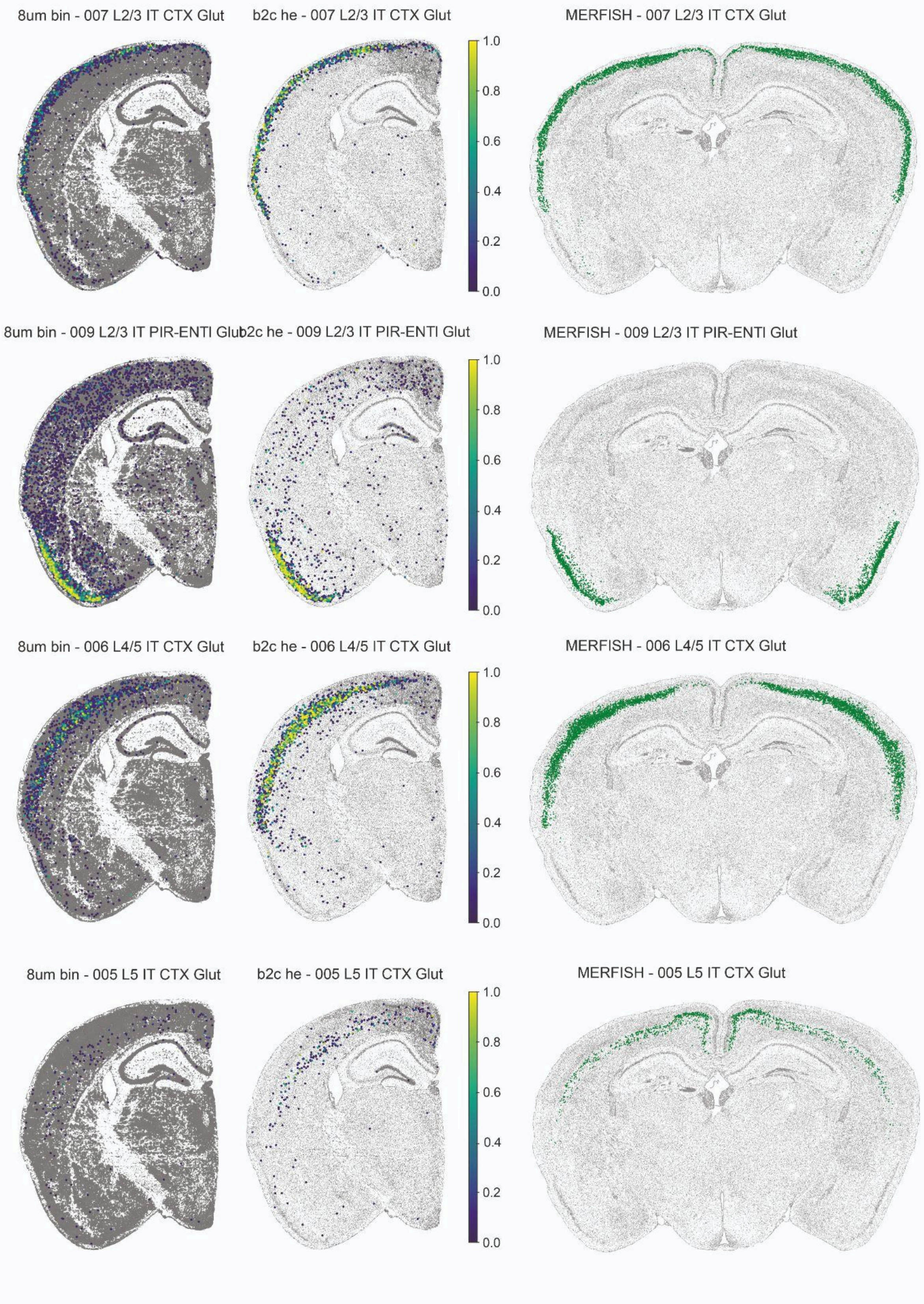
8um bin - 007 L2/3 IT CTX Glut
b2c he - 007 L2/3 IT CTX Glut
MERFISH - 007 L2/3 IT CTX Glut
8um bin - 009 L2/3 IT PIR-ENTI Glut
b2c he - 009 L2/3 IT PIR-ENTI Glut
MERFISH - 009 L2/3 IT PIR-ENTI Glut
8um bin - 006 L4/5 IT CTX Glut
b2c he - 006 L4/5 IT CTX Glut
MERFISH - 006 L4/5 IT CTX Glut
8um bin - 005 L5 IT CTX Glut
b2c he - 005 L5 IT CTX Glut
MERFISH - 005 L5 IT CTX Glut
1.0
0.8
0.6
0.4
0.2
0.0

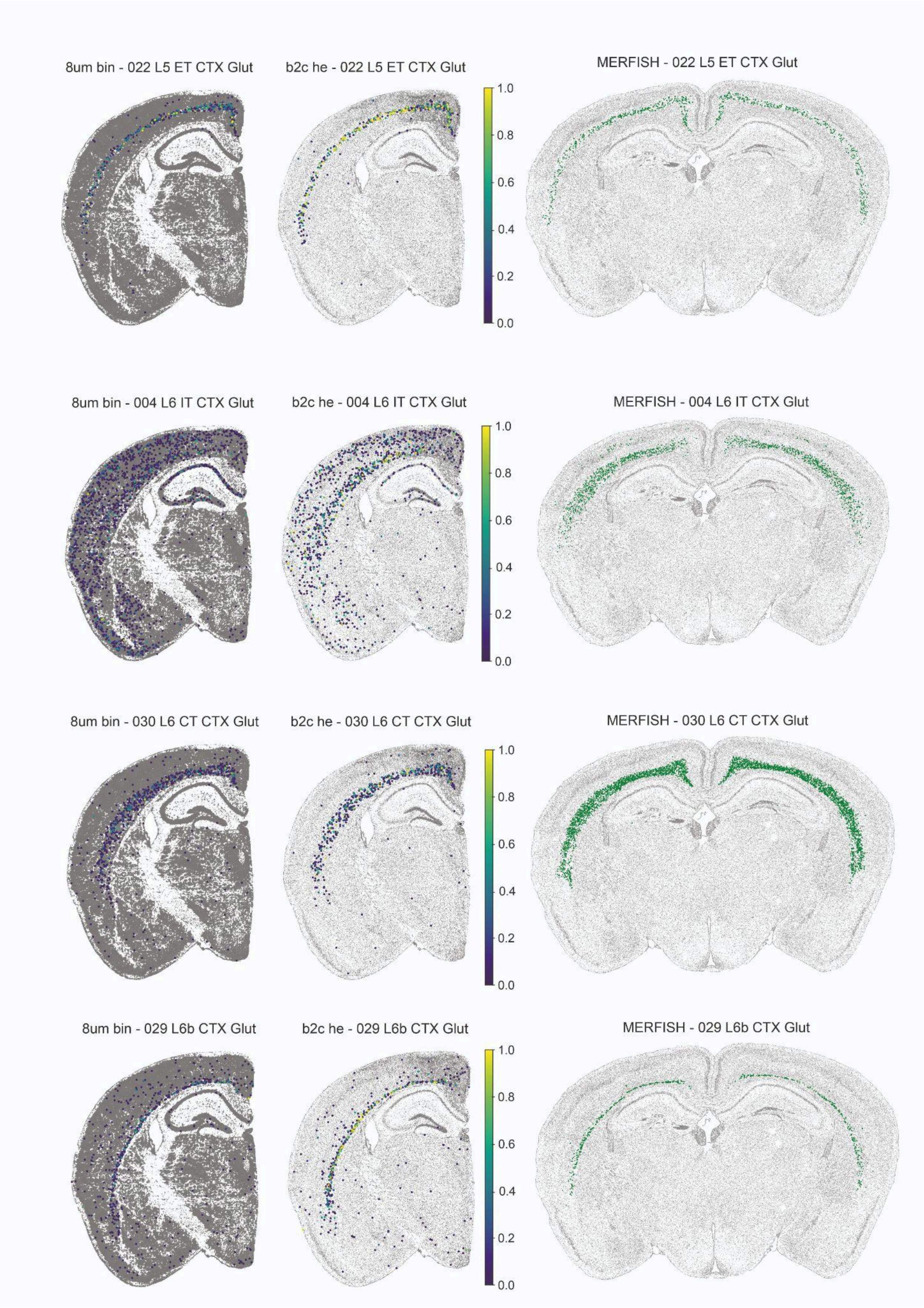

8um bin - 022 L5 ET CTX Glut
b2c he - 022 L5 ET CTX Glut
MERFISH - 022 L5 ET CTX Glut
8um bin - 004 L6 IT CTX Glut
b2c he - 004 L6 IT CTX Glut
MERFISH - 004 L6 IT CTX Glut
8um bin - 030 L6 CT CTX Glut
b2c he - 030 L6 CT CTX Glut
MERFISH - 030 L6 CT CTX Glut
8um bin - 029 L6b CTX Glut
b2c he - 029 L6b CTX Glut
MERFISH - 029 L6b CTX Glut
1.0
0.8
0.6
0.4
0.2
0.0

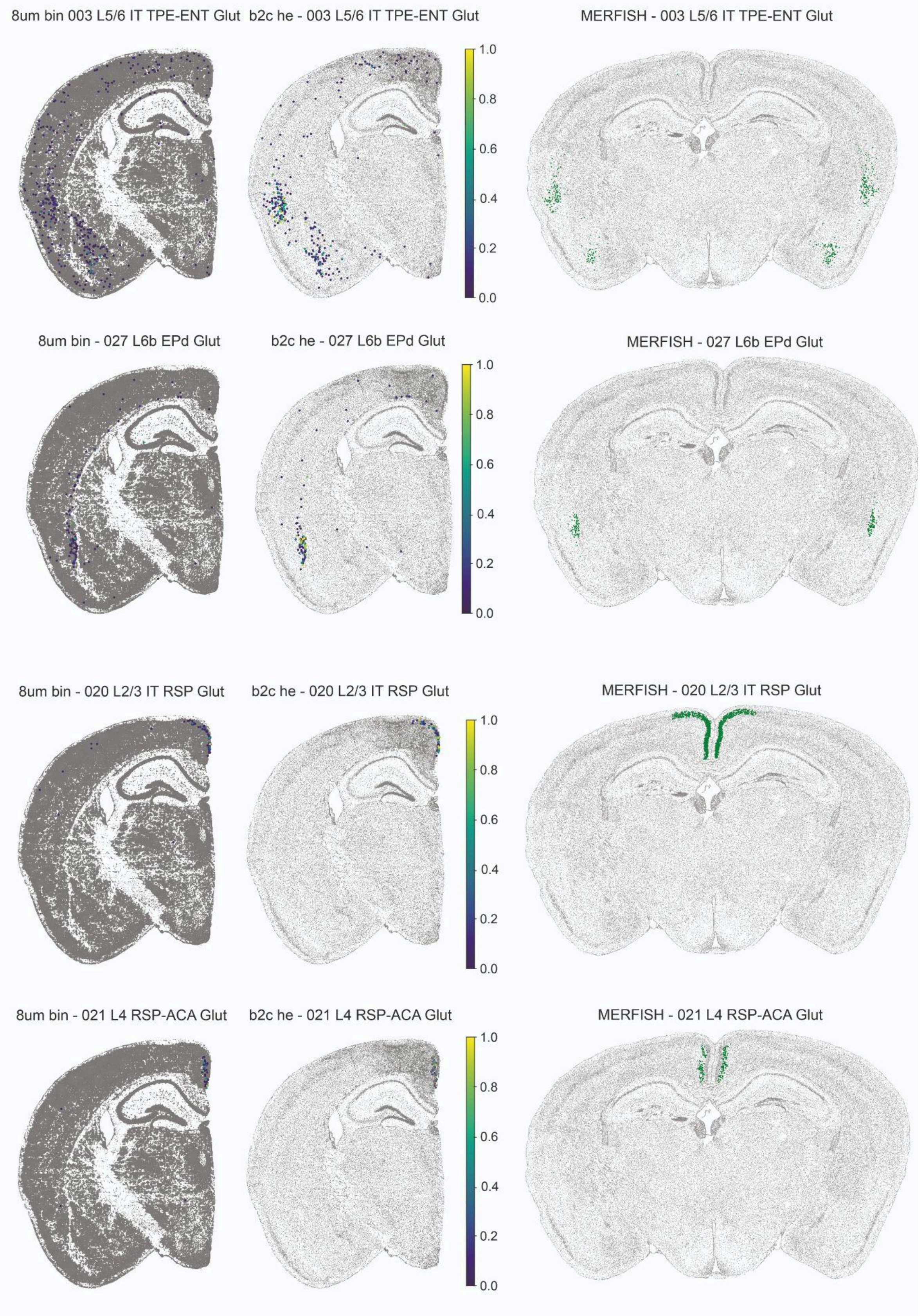
8um bin 003 L5/6 IT TPE-ENT Glut
b2c he - 003 L5/6 IT TPE-ENT Glut
MERFISH - 003 L5/6 IT TPE-ENT Glut
8um bin - 027 L6b EPd Glut
b2c he - 027 L6b EPd Glut
MERFISH - 027 L6b EPd Glut
8um bin - 020 L2/3 IT RSP Glut
b2c he - 020 L2/3 IT RSP Glut
MERFISH - 020 L2/3 IT RSP Glut
8um bin - 021 L4 RSP-ACA Glut
b2c he - 021 L4 RSP-ACA Glut
MERFISH - 021 L4 RSP-ACA Glut
1.0
0.8
0.6
0.4
0.2
0.0

**Supplementary Figures S3-S5. Comparison of individual cortical cell distribution of predicted and reference datasets.** 8μm bins and b2c CellTypist confidence score. MERFISH cell types are represented in green.

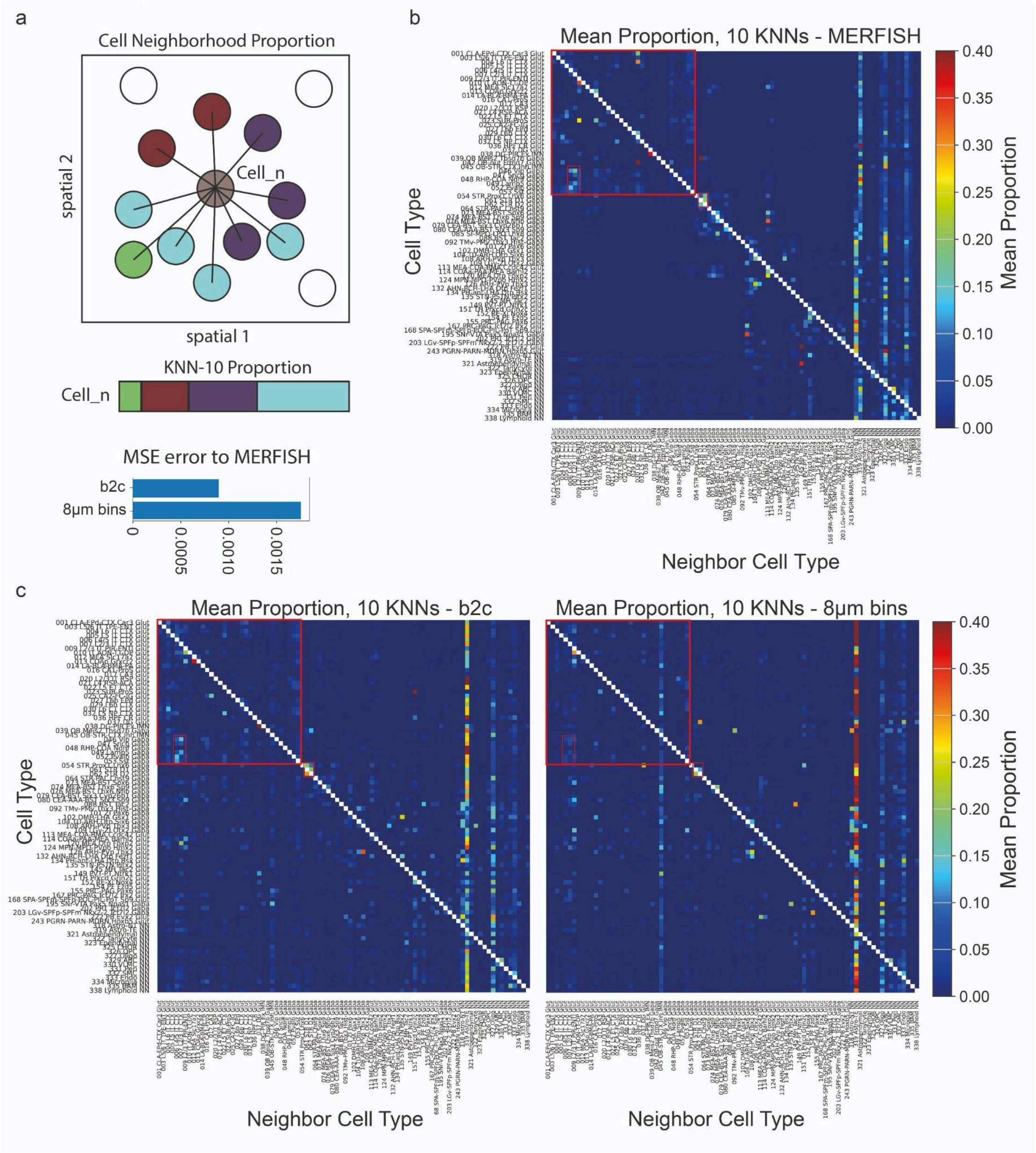


**Supplementary Figure S6. Cell-cell neighbourhood composition evaluation. a.** Illustration of neighbourhood proportion for a given cell to its 10 nearest (euclidean distance) cells from different types (colours)**.** Mean square error of the difference between bin2cell and 8µm bins data for all cells when compared to MERFISH **b.** Mean neighbourhood composition of all cell types for MERFISH data**.** Specific cell associations are highlighted in red. **c.** Mean neighbourhood composition of bin2cell and 8µm bins for predicted cell types (confidence score>0.2)

### Supplementary Section 4: Spatial mapping of human Colorectal cancer

The “Visium HD Spatial Gene Expression Library, Human Colorectal Cancer (FFPE)” Visium HD demo dataset by 10X Genomics was retrieved from https://www.10xgenomics.com/datasets/visium-hd-cytassist-gene-expression-libraries-of-human-crc
A small mismatch between the H&E image and the GEX pattern was corrected by manually adding a factor (-5) to spatial XY coordinates. As part of bin2cell, images were rendered at 0.3 mpp. StarDist H&E segmentation was performed with prob_thresh=0.1 and “fluo” segmentation was performed with prob_thresh=0.01 and nms_thresh=0.1.

#### Cell type prediction with 2 CellTypist models

Since there was no single CellTypist model that matched this dataset, we used a combination of two models: 1) The “Human_Colorectal_Cancer” model from the CellTypist database and a custom model generated from recently curated gut human reference. To reach a unified annotation we then assigned a cell label based on the model that produced the highest confidence score (provided by CellTypist algorithm). Finally, cell types were merged by assigning joint names to similar cells from both models which sometimes diverged in the label format.

#### Prediction performance and gene coverage

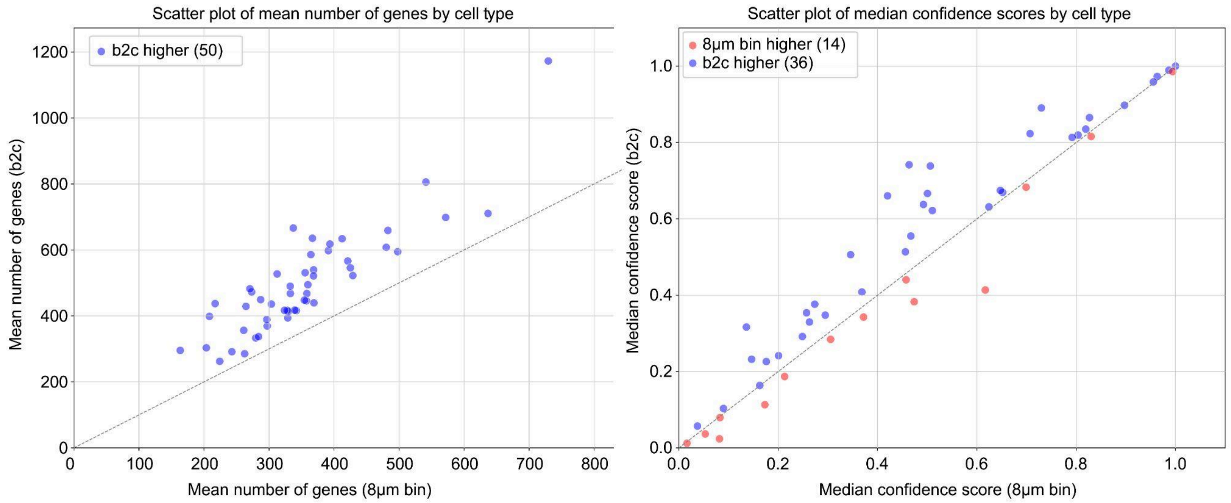


**Supplementary Figure S7.** Mean number of genes in cell types in b2c compared to 8µm bins. CellTypist model median confidence scores for all cell types, a minimum of 0.01 median confidence score for b2c and 8µm bins, blue are cell types that have a higher score for b2c.

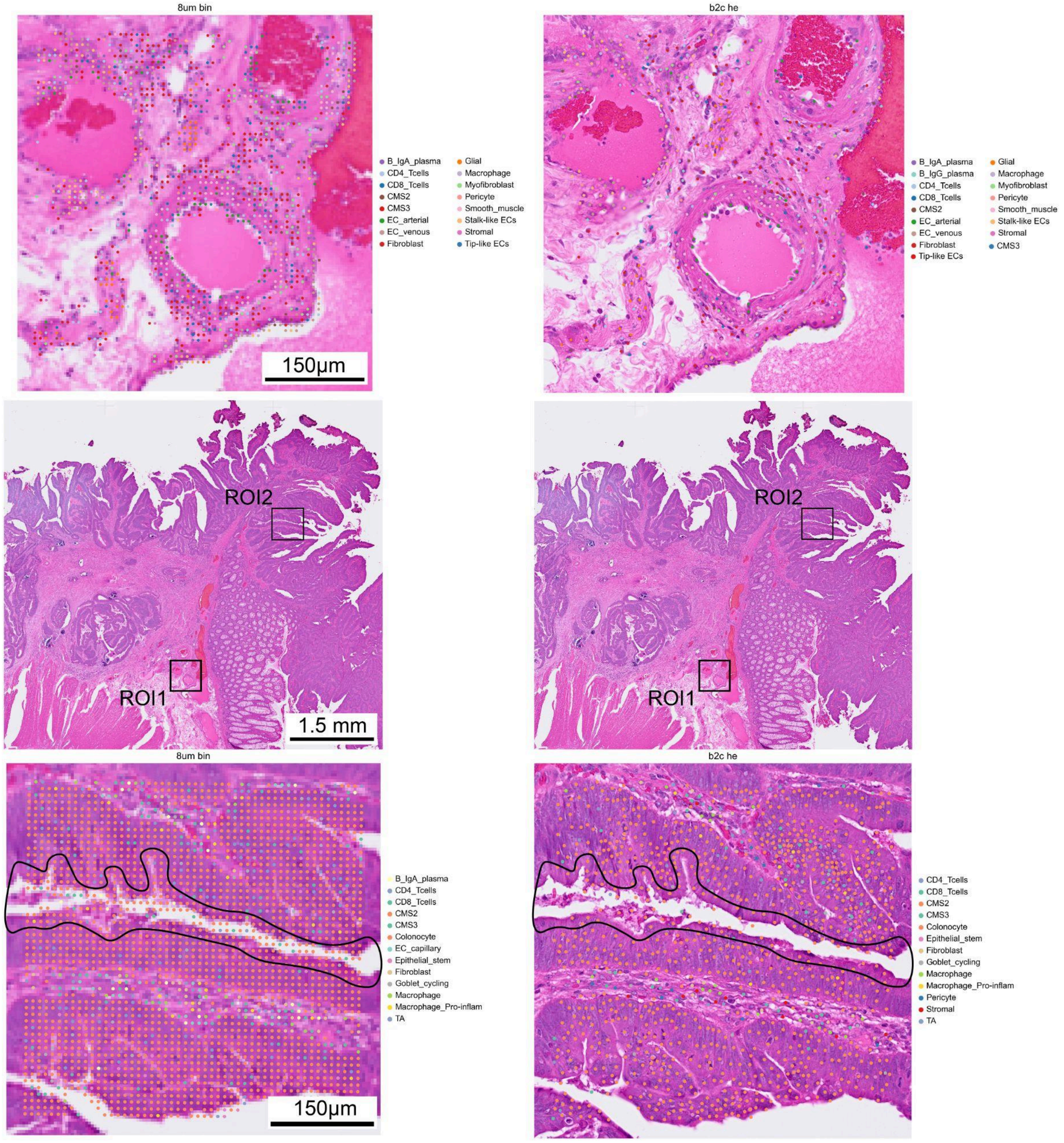


**Supplementary Figure S8. Single cell cell type distribution in vasculature and tissue edges. Top** ROI 1 - magnification of a region with arterial and venous vessels highlighting cellular organisation captured with (Left) 8μm bins and (right) bin2cell. Note that bin2cell captures mono-layered formation of vessel walls and matches the morphology of the H&E image, while 8μm bin predictions are less specific. **Middle.** Overview of the colorectal cancer sample with highlighted regions. **Bottom** ROI 2 - Tissue edges in (left) 8μm bins show non-specific cell predictions in regions which are outside the tissue. Bin2cell is able to accurately group the bins into cells that get predicted as CMS2, the expected cell type in the region.

**Supplementary Section 5: Gene2Probe custom probe design for Visium FFPE and FLEX**

Probe-based spatial transcriptomics technologies offer great flexibility to the user to customise their assays. While technologies such as Visium HD and Visium CytAssist enable the capture of most protein-coding genes, there are many applications that might require the design of custom probes. For example, one might be interested in adding probes for genes not currently included in the panel, such as the female-specific long non-coding RNA *XIST*. Additional applications could include capturing the usage of specific isoforms, such as CD45RA versus CD45RO by different T cell populations, or adding more probes for a gene of interest to maximise its recovery.

Such probes need to fulfil multiple requirements, many of which are often assay- or even version-specific. For example, 10x Genomics currently recommends probes used for Visium HD to be within a specific range of GC content, to have a T in the 25th nucleotide position and to not overlap repeats or common polymorphisms around the ligation junction. Additionally, any probe used to infer transcript abundances needs to be specific to the transcript of interest.

Gene2probe is a flexible pipeline that aims to help the user design such probes. It has been designed mainly around the current guidelines of 10x Genomics for the design of Visium Cytasist and Visium HD probes, but can be easily tuned to accommodate different probe lengths and desired sequence features.

First, the user needs to specify a set of regions of interest, either providing them directly or by specifying the gene ID and feature of interest (e.g., exons, coding sequences, introns or splice junctions). Next, gene2probe will generate all possible k-mers (based on the probe length provided by the user) and use pybedtools (Dale *et al.*, 2011) to intersect them with genomic sets, such as repeats, low complexity regions, gaps in the genome assembly and common polymorphism. K-mers that overlap any of these regions (globally or within a probe subregion specified by the user, such as the ligation junction) are excluded from further analyses. For the remaining regions, gene2probe extracts the DNA sequence and estimates sequence features, such as the GC content of each probe (or probe part for split probes). K-mers fulfilling the requirements set by the user are retained and aligned against the human transcriptome to assess their specificity to the transcript of interest. Gene2probe relies on blastn (Camacho *et al.*, 2009) for this alignment, and includes a manual on how to generate a database for the human transcriptome. Probes with a significant hit in any transcript other than the transcript of interest are discarded, whereas the remaining probes can be prioritised based on the user's preference (for example prioritising shorter homopolymer lengths or a tighter range of GC content). To facilitate the selection of the final probe set, which according to 10x should not overlap each other, gene2probe iteratively selects the top probe from the ranked data frame, then removes all probes within a user-specified window and selects the next probe until the desired number of probes has been identified.

Since probe design can take a lot of fine tuning, with criteria being more lenient or stricter depending on the number of available probes, we implemented gene2probe using a flexible and interactive jupyter notebook interface, which allows the user to easily adjust the requirements of the probe. For example, we recommend users to initially exclude all probes overlapping repeats or common polymorphisms in any position. However, if the number of resulting probes is too small (e.g., due to a short region of interest), the user can relax this requirement, only removing probes with overlaps within a certain range (e.g., +/-5 nucleotides from the ligation junction). Furthermore, this flexibility in parameter choice allows gene2probe to be used for the design of probes for different assays.

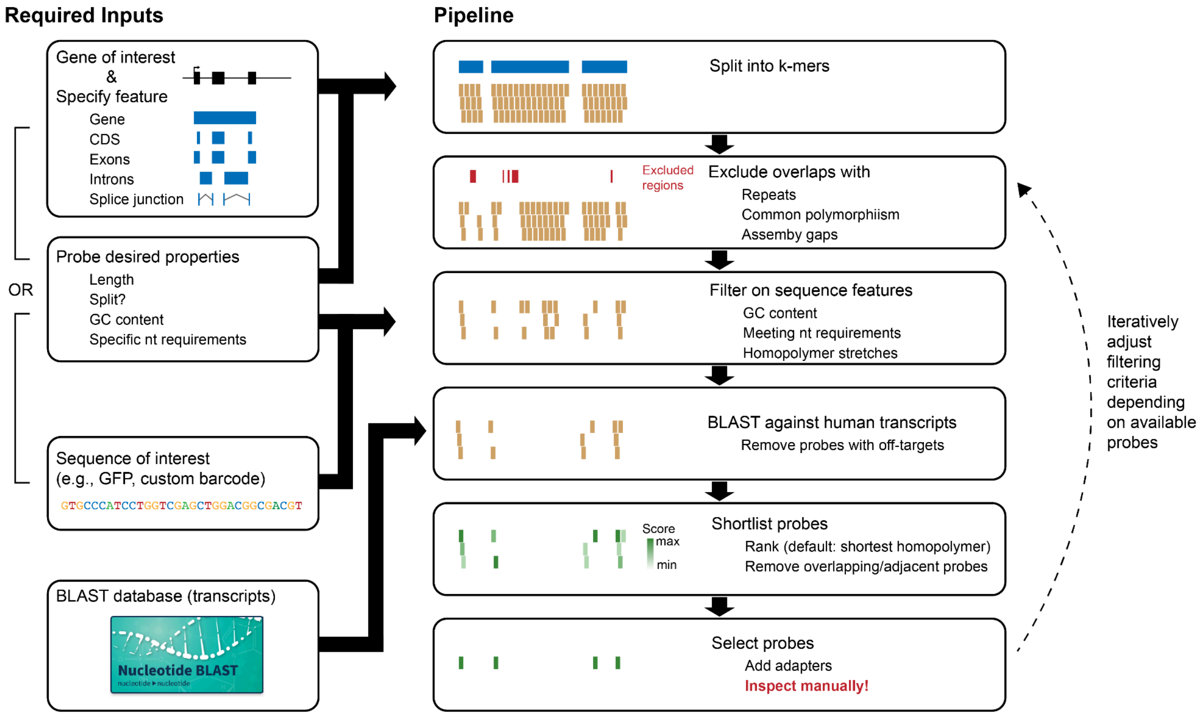


**Supplementary Figure S9. Schematic overview of the gene2probe pipeline.** Given a gene or sequence of interest and a set of desired probe properties, gene2probe will split the sequence into all possible k-mers for a probe of length k. It will then exclude k-mers that overlap undesirable regions, such as repeats and common polymorphisms, and k-mers that do not satisfy the sequence requirements, such as falling outside the specified GC content range. Gene2probe then uses BLAST to investigate matches of the k-mers to all human transcripts and excludes k-mers with potential off-targets. Finally, the user can rank probes based on custom criteria and select a non-overlapping set of highly scoring probes. The precise filtering criteria can be adjusted depending on the number of probes that are available for the gene or sequence of interest.
